# Comparative impact of porcine reproductive and respiratory virus and swine influenza A virus infections on respiratory lymph nodes B cells and macrophages

**DOI:** 10.1101/2024.10.11.617816

**Authors:** C. Hervet, A. Perrin, P. Renson, C. Deblanc, M. Muñoz, F. Meurens, J. Argilaguet, G. Simon, O. Bourry, P. Maisonnasse, N. Bertho

**Affiliations:** Oniris, INRAE, BIOEPAR, 44300, Nantes, France; Swine Virology Immunology Unit, Ploufragan-Plouzané-Niort Laboratory, French Agency for Food, Environmental and Occupational Health and Safety (ANSES), 22440, Ploufragan, France; Unitat Mixta d’Investigació IRTA-UAB en Sanitat Animal, Centre de Recerca en Sanitat Animal (CReSA), Campus de la Universitat Autònoma de Barcelona (UAB), 08193 Bellaterra, Spain; Institut de Recerca i Tecnologia Agroalimentàries (IRTA), Programa de Sanitat Animal, Centre de Recerca en Sanitat Animal (CReSA), Campus de la Universitat Autònoma de Barcelona (UAB), 08193 Bellaterra, Spain; WOAH Collaborating Centre for the Research and Control of Emerging and Re-Emerging Swine Diseases in Europe (IRTA-CReSA), 08193 Bellaterra, Spain; Department of Veterinary Microbiology and Immunology, Western College of Veterinary Medicine, University of Saskatchewan, Saskatoon, SK, Canada; CRIPA, Fonds de Recherche du Québec, Département de pathologie et microbiologie, Faculté de médecine vétérinaire, Université de Montréal, Saint-Hyacinthe, QC, Canada

## Abstract

Porcine Reproductive and Respiratory Syndrome Virus (PRRSV) strongly impacts the pig rearing sector due to its persistence in infected animals. Interestingly, although the PRRSV family exhibits considerable genome variability, with the PRRSV-1 and PRRSV-2 subtypes having been finally classified in two different species (Betaarterivirus suid 1 and 2). Both viruses, as well as their derived-attenuated vaccine strains, persist for months, due in part to their ability to delay the appearance of neutralizing antibodies. Thanks to extensive efforts over the past years, we have developed the capability to perform in-depth analysis of the previously poorly understood porcine inverted lymph node (LN). In this study, by comparing the early stages of LN B cell maturation upon PRRSV-1 infection to those induced upon the acute swine influenza A virus infection, we highlighted PRRSV-specific mechanisms, including the expression of PD-L1 in efferent macrophages, the induction of extrafollicular plasmocytes, and the influx of inflammatory monocytes/macrophages. Studies on PRRSV-2 infections report observations compatible with our results, that thus might be generalized to all PRRSV-strains. Moreover, these mechanisms can be compared with those used by the human immunodeficiency virus (HIV) and the murine chronic lymphocytic choriomeningitis virus (LCMV) to hijack the immune response. These similarities can be harnessed to develop new strategies to improve the development of more efficient anti-PRRSV vaccines.

## Introduction

Porcine Reproductive and Respiratory Syndrome Virus (PRRSV) is an enveloped positive-stranded RNA virus from the *Arteriviridae* family with a tropism for macrophages (MΦ). PRRSV can be divided into two species, Betaarterivirus suid 1 (PRRSV-1) and Betaarterivirus suid 2 (PRRSV-2), which share 60% nucleotide identity (1). The virus causes reproductive disorders in sows and respiratory diseases in piglets. Moreover, the virus’ persistence facilitates respiratory super-infections, leading to an important economic impact (2, 3) and an important issue from a One Health perspective (4). Moreover, some emerging American, Chinese, and European (such as Lena) strains are highly pathogenic and threaten the West European pig industry (5–7). Highly pathogenic strains are characterized by high viral loads, severe general clinical signs and high mortality, in sows, weaners and growers.

PRRSV persistence mechanisms are still debated. Among the different PRRSV strains from PRRSV-1 and -2 species, some induce strong Th1 response (PRRSV 1.2 Lena), some induce very little T cell response (PRRSV 1.1), and others induce a strong Treg response (PRRSV-2) (8–10). This heterogeneity in T helper induction among PRRSV strains suggests that T cell responses are not critical for PRRSV persistence in newly infected adult animals. PRRSV-mediated thymus T cells differentiation perturbations have been demonstrated in newborn piglets (11, 12), which can explain the much stronger impact of PRRSV infection on young animals. However, this can hardly be generalized to adult pigs, which are much less dependent on thymic emigrants to mount an efficient immune response. All PRRSV strains delay the appearance of neutralizing antibodies (NAb) up to one-month post-infection (pi)(13), in both newborn and adult pigs, whereas the non-NAb production occurs within the expected time frame of one-week pi (13). By comparison, swine influenza A virus (swIAV), another major porcine respiratory pathogen, evokes NAb by one-week pi (14). Interestingly, it has been previously observed that unlike swIAV, PRRSV infection using PRRSV-2 and germfree piglets (15) triggers proliferation of germline-encoded B cells of all isotypes, bearing hydrophobic heavy chain CDR3, with little evidence of immunoglobulin diversification (16, 17). It has been observed that the virus, while no longer present in the lung, can be detected in lymph nodes (LN) for months pi. These data led us to interrogate the impact of PRRSV on the B cell maturation process in the very place where it occurs: the LN.

To fulfill this task, we first had to better understand the peculiar structure of the porcine inverted LN (18). In a previous work (Bordet et al., 2019), we set up flow cytometry gating and sorting strategies that allowed us to define three LN MΦ populations, to which we tentatively assigned mouse counterparts. The perifollicular MΦ (pfMΦ) are positioned in contact with B cell follicles and have a CD163^neg^/CD169^pos^ phenotype similar to the murine subcapsular sinus MΦ. A CD163^pos^/CD169^neg^ MΦ population which we named cord MΦ due to their phenotypic similarity with murine cord MΦ. They are positioned in what we initially identified as the LN cord. It later appeared that these CD163^pos^/CD169^neg^ cells were localized in the trabecula (the collagen structure supporting the lymphatic sinuses), the lymphatic sinus itself and the T cell area, probably representing different MΦ types. For better accuracy, herein we will name them CD163^pos^/CD169^neg^ MΦ. The last MΦ population is CD163^pos^/CD169^pos^ is positioned in the LN parenchyma, at the periphery of the LN corresponding, in inverted LN, to the LN exit. We therefor named it efferent MΦ (effMΦ). We also set protocols to follow B cells positioning and maturation process (Bordet et al., 2019).

In the present work, we used this toolbox to compare the early divergences between anti-PRRSV and anti-swIAV innate and adaptive immune responses in the tracheobronchial LN. We opted for an in-depth LN analysis, which required the culling of the animals, but prevented the possibility to conduct a time course study. We have therefore chosen to analyze the LN at 8 dpi, when the most contrasted anti-swIAV and anti-PRRSV immune responses might be expected. At this time, a well-established anti-swIAV immune response can be anticipated, with the appearance of the first NAb. In contrast, the PRRSV infection might have had the time to set up its mechanisms of suppression of the immune response.

## Material and Methods

### Virus Production and Titration

The highly pathogenic Lena PRRSV-1.3 strain was kindly provided by Dr. Hans Nauwynck, (University of Ghent, Belgium). Lena has been isolated in Belarus in 2007 from a herd with mortality, reproductive failures and respiratory disorders (7). The Lena viral stock for *in vivo* experiment was produced using fresh primary alveolar macrophages (AM) collected from adult animals from the controlled-PRRSV free INRAE herd of the Unité Expérimentale de Physiologie Animale de l’Orfrasière (UEPAO, INRAE, Nouzilly, France). Supernatants from infected cells were clarified by centrifugation at 3,300g, filtered on 0.8µm. Virus titration was performed on fresh primary AM using the Ramakrishnan TCID50 method (21).

The european human-like reassortant swine H1N2 (swIAV H1N2) strain A/swine/Ille-et-Vilaine/0415/2011 was selected among the collection of the French National Reference Laboratory for Swine Influenza (ANSES, Ploufragan, France). It was isolated from a nasal swab taken from a pig with acute respiratory disease in a herd located in Brittany, France. It was propagated on Madin-Darby Canine Kidney (MDCK) (ATCC reference CCL-34) cells for 24 hours (h) in DMEM medium supplemented with 10% Fetal calf serum, 1% of Streptomycin/Penicillin/Amphotericin solution (Eurobio scientific, Les Ulis, France), and 2 µg/mL of trypsin TPCK (Worthington Biochemical Corp., Lakewood, NJ, USA). The supernatant was then collected, clarified by centrifugation (600× g, 20 min) and stored at −80°C.

Both virus stocks were concentrated and purified on Amicon Ultra-15 centrifugal Filters (Sigma-Aldrich – reference number UFC910024 – pore size 100 kDa Nominal Molecular Weight cut-off) after a 20 min centrifugation at 4000× g and 4⍰C. Titer determinations of swIAV H1N2 were carried out on MDCK using Reed & Muench method (22). The viral titers of purified swIAV H1N2 reached 9.8.10^6^ TCID50/mL, while PRRSV-1 stock titer was 2.10^9^ TCID50/mL.

### Animals, *in vivo* infections and tissue collection

Infection experiments were conducted at CReSA (Barcelona, Spain). The treatment, housing, and husbandry conditions conformed to the European Union Guidelines (Directive 2010/63/ EU on the protection of animals used for scientific purposes). Animal care and procedures were in accordance with the guidelines of the Good Laboratory Practices under the supervision of the Ethical and Animal Welfare Committee of the Universitat Autonoma de Barcelona (number 1189) and under the supervision of the Ethical and Animal Welfare Committee of the Government of Catalonia (number 5796). Twenty pigs (7–8 weeks old, LandracexPietrain) were housed in separate isolation rooms and were randomly assigned to three experimental groups of six (Mock and swIAV-H1N2) or eight (PRRSV-Lena) pigs. The animals were seronegative to IAV (ID Screen Influenza A Antibody Competition ELISA, ID-Vet, Grabels, France) and PRRSV (IDEXX PRRS X3 5/S TRIP), and free of swIAV and PRRSV current infection at the start of the experiment, as tested by PCR using qPCR protocols described in (23) for swIAV and RT-qPCR Kit from Applied biosystems (A35751 VETMAX PRRSV DETECTION KIT 100 RXN - Thermo Fisher) for PRRSV. On day 0 (after one week of acclimatization period), 11 weeks oldanimals were intranasally infected by 1.10^6^ TCID50 swIAV-H1N2 (5.10^5^/ml per nostril) or 1.10^6^ TCID50 Lena (5.10^5^/ml per nostril). Pigs were examined daily; clinical score was the aggregation of four criterions scored from 0 to 3 (**Suppl. Table 1**). Three of them were monitored daily (behavior, respiratory signs, rectal temperature) and one twice a week (body weight). Nasal swabs were collected daily from 0 to 8 dpi in 500 µl Roswell Park Memorial Institute (RPMI) medium and immediately frozen at −80°C. Sera were collected at 0 and 8 dpi and store at -20°C.

### Anti-swIAV and anti-PRRSV IgG, IgM and NAb detection in sera

The immunoglobulins G (IgG) directed against the swIAV were quantified with the ID Screen® Influenza A Antibody Competition Multispecies kit (Innovative Diagnostics, Grabels, France) following manufacturer’s instructions. IgG levels were expressed in inhibition percentage.

Anti-swIAV immunoglobulins M (IgM) were detected with the kit ID Screen® Influenza A Nucleoprotein Swine Indirect kit (Innovative Diagnostics, Grabels, France) with a modified protocol using in-house controls and a goat anti-pig IgM antibody HRP conjugate (A100-117, Bethyl—Fortis Life Sciences, Montgomery, TX, USA) at a 1:10000 dilution as a conjugated antibody. IgM levels were expressed in sample-to-positive (S/P) ratios.

NAb targeting the H1N2 strain were quantified by a virus neutralization assay as previously described (Deblanc et al, 2020). The NAb titer was determined as the reciprocal of the highest dilution of serum that prevents virus infection of the cell monolayer, determined by the absence of cytopathic effect. A serum was considered positive when virus neutralization titer ≥ 20.

Levels of immunoglobulins G (IgG) directed against PRRSV (N protein) were assessed using the Pigtype PRRS Ab ELISA kit (Indical Bioscience GmbH, Leipzig, Germany) following manufacturer’s instructions and expressed as sample-to-positive (S/P) ratios. Anti-PRRSV (N protein) IgM levels were measured using the same PRRSV ELISA kit, replacing the kit’s conjugated antibody by a goat anti-pig IgM HRP-conjugated antibody (A100-117, Bethyl Laboratories—Fortis Life Sciences, Montgomery, TX, USA) at 1:25000 dilution and replacing the commercial controls by in-house calibrated negative and positive serum controls to calculate sample-to-positive (S/P) ratios. PRRSV-specific NAb targeting the Lena strain were quantified in serum using MARC145 cells. Heat-inactivated sera were two-fold serially diluted at 56 °C for 30 min, and then 50 µl of each dilution was incubated in duplicate in 96-well microtiter plates with the MLV1 DV strain at 101 ± 0.5 TCID50/50 µl for 1 h at 37°C with rocking agitation. A suspension of MARC-145 cells (0.5⍰×⍰105 per well) was then added. After incubation for five days at 37 °C, the titers were determined as the reciprocal of the highest dilution of serum that prevented virus infection of the cell monolayer, as determined by the absence of cytopathic effects in half of the duplicate wells. A serum was considered positive when virus neutralization titre ≥ 10.

### Serum cytokines dosage

Serum cytokines concentrations were determined according to manufacturer instructions for CXCL13, IL-5, BAFF, IL-2, IL-4, IL-6, IL-13 IL-17 and IL-10 and IFN-γ. CXCL13, IL-5 and CD257 were quantified by sandwich ELISA with kits from LSBio (LS-F41718 Porcine BLC/CXCL13 and LS-F45145 Porcine IL-5) or from Biosource (MBS2512446 Porcine BAFF/CD257). Other cytokines were measured by 2-plex (Porcine IL-10, IFN-γ) or 5-plex (Porcine IL-4, IL-13, IL-17, IL-6, IL-2) Luminex assays. All kits were purchased to Abyntek Biopharma S.L. (Bizkaia, Spain).

### Lymph node collection and cells isolation

At 8 dpi, animals were culled and lungs were collected. The tracheobronchial LN in the precisely delimited space between the right main bronchia and the accessory right bronchia were collected for dissociation. LN were minced and incubated with RPMI 1640 supplemented with 100 IU/ml penicillin, 100 mg/ml streptomycin, 250 ng/ml Fungizone® (Antibiotic-Antimycotic 100X ThermoFisher Scientific, Illkirch, France), 2mM L-glutamine and 10% heat inactivated fetal calf serum (FCS, Invitrogen, Paisley, UK). Digestion was performed for 30 min at 37⍰C with 2 mg/ml collagenase D (Roche, Meylan, France), 1 mg/ml dispase (Invitrogen) and 0.1 mg/ml DNase I (Roche). Filtration on 40µm cell strainers was performed and red blood cells were lysed using erythrolysis buffer (10mM NaHCO3, 155mM NH4Cl, and 10mM EDTA). Cells were washed in PBS/EDTA, and re-suspended in FCS + 10% dimethyl sulfoxide (DMSO) for liquid nitrogen storage before cell sorting.

The tracheobronchial LN situated at the bifurcation of the two main bronchia were collected for Tissue Teck (Sakura, Paris, France) embedding and were frozen on dry ice before -80°C storage.

### Flow Cytometry Analysis and Cell Sorting

Cells were stained in a blocking solution, composed of PBS-EDTA (2mM) supplemented with 5% horse serum and 5% swine serum. Staining was made in 4 steps, including PBS/EDTA with 2% FCS washing between each step: uncoupled primary anti-CD169, anti-IgM and anti-CD172a antibodies (Table 1) were added to the blocking solution for 30min on ice and then washed. Fluorescent, secondary, mouse isotype specific antibodies were then added, respectively anti-IgG2a-PE-Cy7, anti-IgM-Dye-Light-649 and anti-IgG2b-APC-Cy7 (Table 1) for 20 min on ice and then washed. Then, to saturate the potential unbound IgG1-directed secondary antibodies sites, cells were incubated for 30 min in the blocking solution supplemented with isotype-control IgG1 (10 µg/ml). Third, the fluorochrome-coupled primary IgG1: anti-CD21 coupled to FITC, anti-CD163 coupled to PE (Table 1) were added for 30 min on ice and then washed before re-suspension in a DAPI-containing buffer for sorting. Sorting was performed using a BD FACSAria II cell sorter (Becton Dickinson, San Jose, USA) equipped with 407, 488, and 633nm lasers. Temperature was kept at 4⍰C during the whole sorting. FlowJo software (version X.1.0, Tree Star, Ashland, OR, USA) was used for analysis. Sorted cells were centrifuged and lysed in RLT buffer for storage at -80°C before further processing.

**Table 1:** Antibodies used for flow cytometry and fluorescent microscopy analysis.

| Ab | Clone | Species/Isotype | Fluorophore | Supplier |
| --- | --- | --- | --- | --- |
| **Primary Abs** |  |  |  |  |
| anti-CD169 | 1F1 CR4 | mIgG2a | none | J. Dominguez, INIA |
| anti-CD163 | 2A100 | mIgG1 | PE | Serotec |
| anti-CD163 | 2A100 | mIgG1 | none | Serotec |
| anti-CD21 | B-ly4 | mIgG1 | FITC | BD-Pharma |
| anti-CD21 | B-ly4 | mIgG1 | none | BD-Pharma |
| anti-IgM | PG145A | mIgM | none | MAC-WSU |
| anti-CD172a | 74-22-15a | mIgG2b | none | MAC-WSU |
| anti-Pax5 | 1H9 | rat | none | Thermofisher |
| anti-Bcl6 | K112-91 | mIgG1 | none | BD-Pharma |
| **Secondary Abs** |  |  |  |  |
| anti-mIgM |  | goat | Alexa-647 | Invitrogen |
| anti-mIgG2b |  | goat | APC-Cy7 | Invitrogen |
| anti-mIgG2a |  | goat | PE-Cy7 | Invitrogen |
| anti-mIgG2a |  | goat | Alexa-647 | Invitrogen |
| anti-rIgG |  | goat | Alexa-488 | Invitrogen |
| anti-rIgG |  | goat | Alexa-647 | Invitrogen |
| anti-mIgG1 |  | goat | Alexa-555 | Invitrogen |

### RT-qPCR

Total RNA in RLT Buffer were extracted using RNeasy MicroKit (Qiagen) following the manufacturer’s instructions. Total RNA quantity and quality were assessed using Nanophotometer (Implen, Munich, Germany). cDNA were generated with a reverse trancriptase in the iScript Reverse Transcription Supermix for RT-qPCR (Bio-Rad, Hercules, CA, USA). The generated cDNAs were then diluted (2×) and combined with the primer set and SYBR Green Supermix (Bio-Rad) following the manufacturer’s recommendations. PRRSV-Lena and swIAV genomes were detected by multiplex qPCR (Eurogentec, Liège, Belgium) with probes and primers specified in **Table 2**.

**Table 2:**
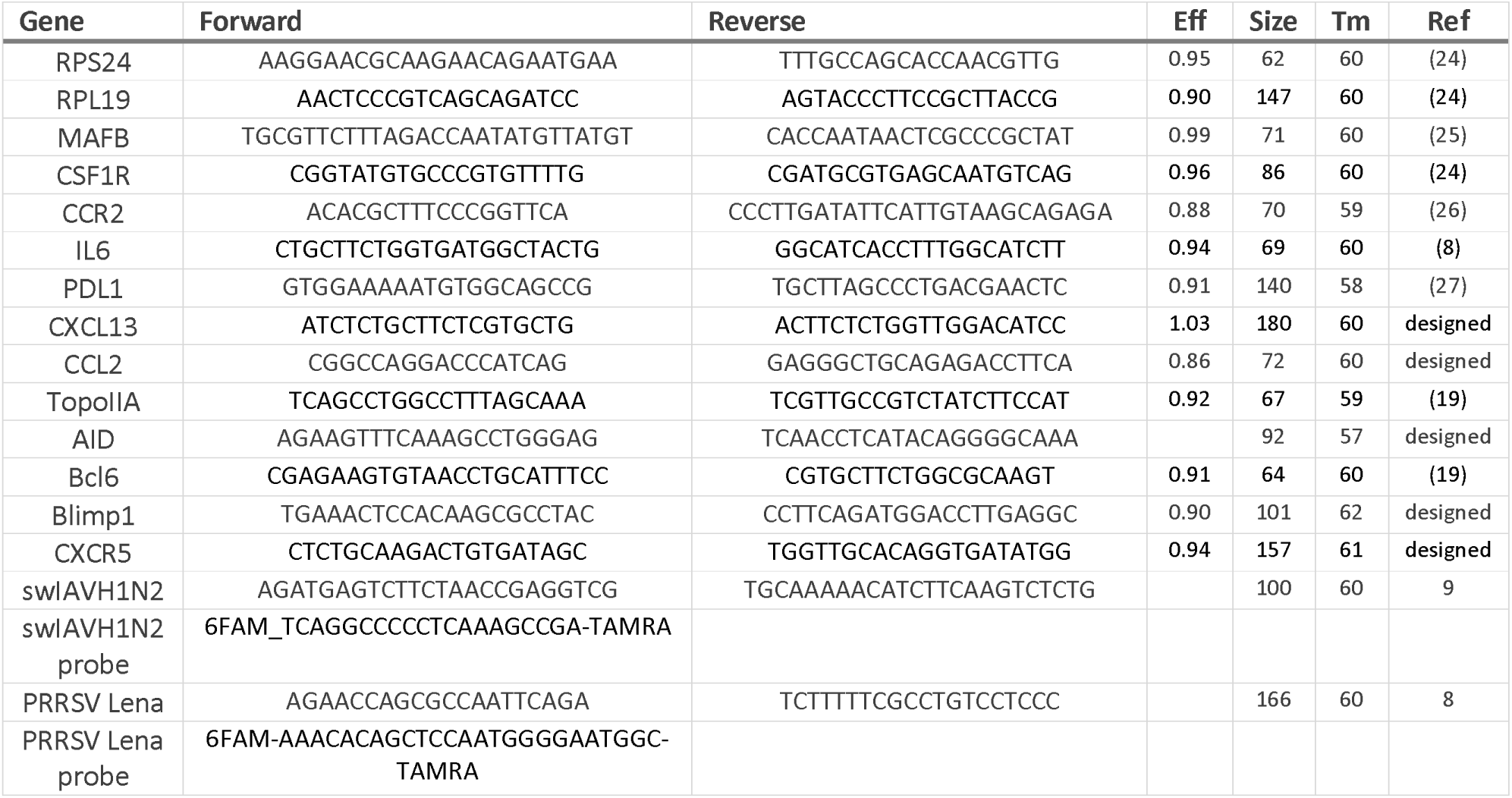
RT-qPCR primers.

### Immunohistochemical staining

Sections of 14 µm were obtained from Tissue Teck embedded LN using a cryostat (Leica CM3050S, Nanterre, France) and deposed on Superfrost® glass slides (ThermoFisher scientific). Cryosections were fixed in methanol/acetone (1:1) at −20⍰C for 20 min. Fixed slides were saturated using PBS supplemented with 5% horse serum and 5% swine serum for 30 min at room temperature (RT). Primary and secondary antibodies (**Table 1**) were added at 4⍰C overnight or for 30 min, respectively. Acquisition was performed using the same settings for all slides to ensure comparable fluorescence parameters. Microscopy cell counting was performed using Zen software (Zeiss). For intra- and interfollicular centroblasts (CB) and centrocytes (CC) numbering, a virtual line was drawn across the follicle, extending 20% out of the follicle’s diameter on each side. The follicular area limit was considered reached when two CD21 pixels showed a relative fluorescence above 500. Cells were identified along this line as signals having at least 3 pixels above 250 (for Alexa-555) or 550 (for Alexa-647) of relative fluorescence, representing respectively BcL6 and Pax5 expressions. LN from 4 different animals from the 3 infectious conditions (Mock, PRRSV, swIAV) were analyzed. Two or three follicles per LN were blindly counted.

For MΦ numbering, T cell areas were defined as areas between two Pax5-positive follicles and trabeculae were defined as areas with rare DAPI staining along Pax5-positive follicles. LN from 3 different animals from the 3 infectious conditions (Mock, PRRSV, swIAV) were analyzed. Three different T cell areas per LN were blindly counted. A fluorescent signal in one of the 2 channels defining MΦ (Alexa-555 and Alexa-647, defining respectively CD163 and CD169 expressions) with a relative fluorescence above 300 and composed of at least 3 pixels, was counted as one cell. For both MΦ and B cells staining, in some areas, it was impossible to distinguish individual cells within dense cell clusters. In such case, a continuous signal exceeding 10 µm was considered to represent 2 cells, a signal above 20 µm represented 3 cells, and so on.

### Statistics

Graph Pad Prism 10.2.3 (GraphPad Software) was used for graph design and statistics. Shapiro-Wilk test was used to assess the distribution of the data. Between infectious conditions, an unpaired non-parametric Mann-Whitney test was used. When intra-infectious conditions were compared (0 dpi *vs* 8 dpi or different B cells or MΦ populations), a paired non-parametric Wilcoxon rank test was made.

## Results

### PRRSV Lena triggered higher viral load and clinical signs than swIAV, associated with a lower lymph node cellularity and the absence of neutralizing antibodies

Seven/eight weeks old animals were intranasally infected with a dose of 1.10^6^ TCID50 of swIAV-H1N2 or PRRSV-1 Lena. Strong swIAV shedding was observed at 2 dpi before decreasing. Influenza virus was no longer detected in nasal swabs by 8 dpi (**Fig. 1A**). Conversely, PRRSV shedding appeared at 2 dpi similarly to swIAV. However, PRRSV levels continued to increase at 4 dpi before gradually decreasing by 8 dpi (**Fig. 1A**). As expected, intranasal swIAV infection triggered few clinical signs, that were totally resolved at 6 dpi, whereas the pathogenic Lena PRRSV 1.3 strain triggered clear clinical signs for the 8 infected animals at 3 dpi, with no improvement at 8 dpi (**Fig. 1B**).

**Figure 1:**
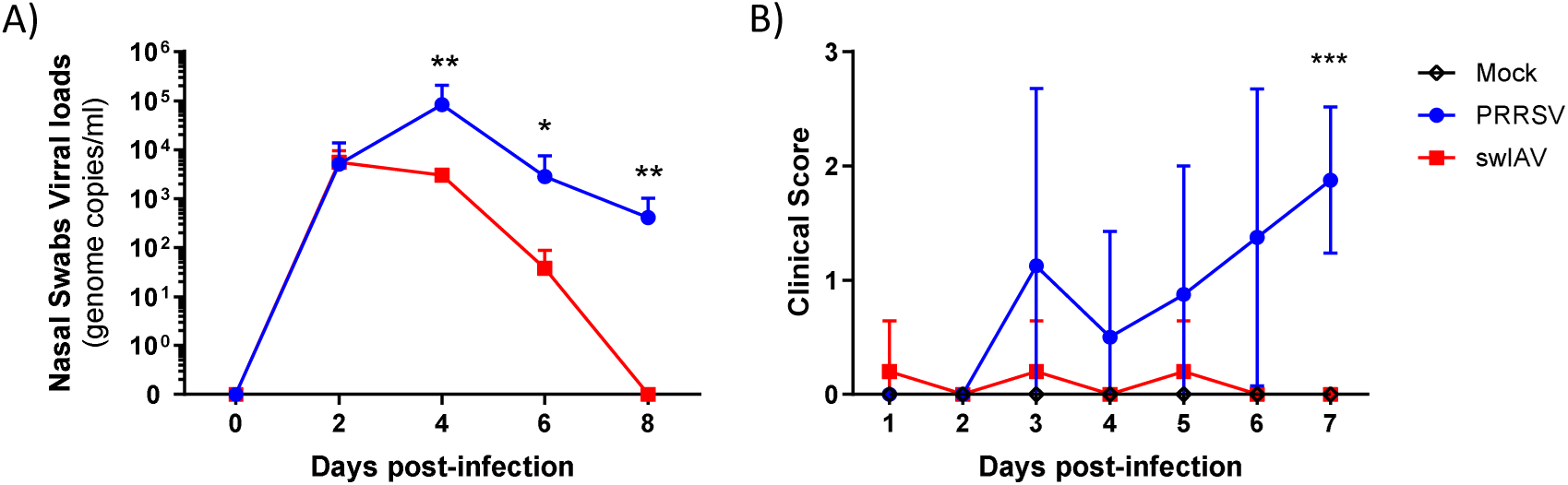
PRRSV Lena and swIAV H1N2 infected pigs present contrasting viral loads and clinical signs. Pigs were intranasally exposed to either PBS (Mock - black - n=6), 1.10^6^ TCID50 of a PRRSV Lena strain (PRRSV - blue - n=8) or 1.10^6^ TCID50 of a swIAV H1N2 strain (swIAV - red - n=6). A) RNA viral loads measured by RT-qPCR in nasal swabs (means with SD). B) Clinical scores (mean with range) were recorded from day 1 to day 7 post-infection. Mann-Whitney unpaired two-tailed t-test, *p* values: *<0.05, **<0.01, ***0.001.

We then validated the humoral immune response course upon viral infections by measuring serum anti-swIAV and anti-PRRSV IgM, IgG and NAb titers. The 3 types of antibodies directed against swIAV were readily detected at 8 dpi (**Fig. 2A**). Conversely, whereas anti-PRRSV IgM and IgG serum antibodies were present, no anti-PRRSV NAb were observed (**Fig. 2B**).

**Figure 2:**
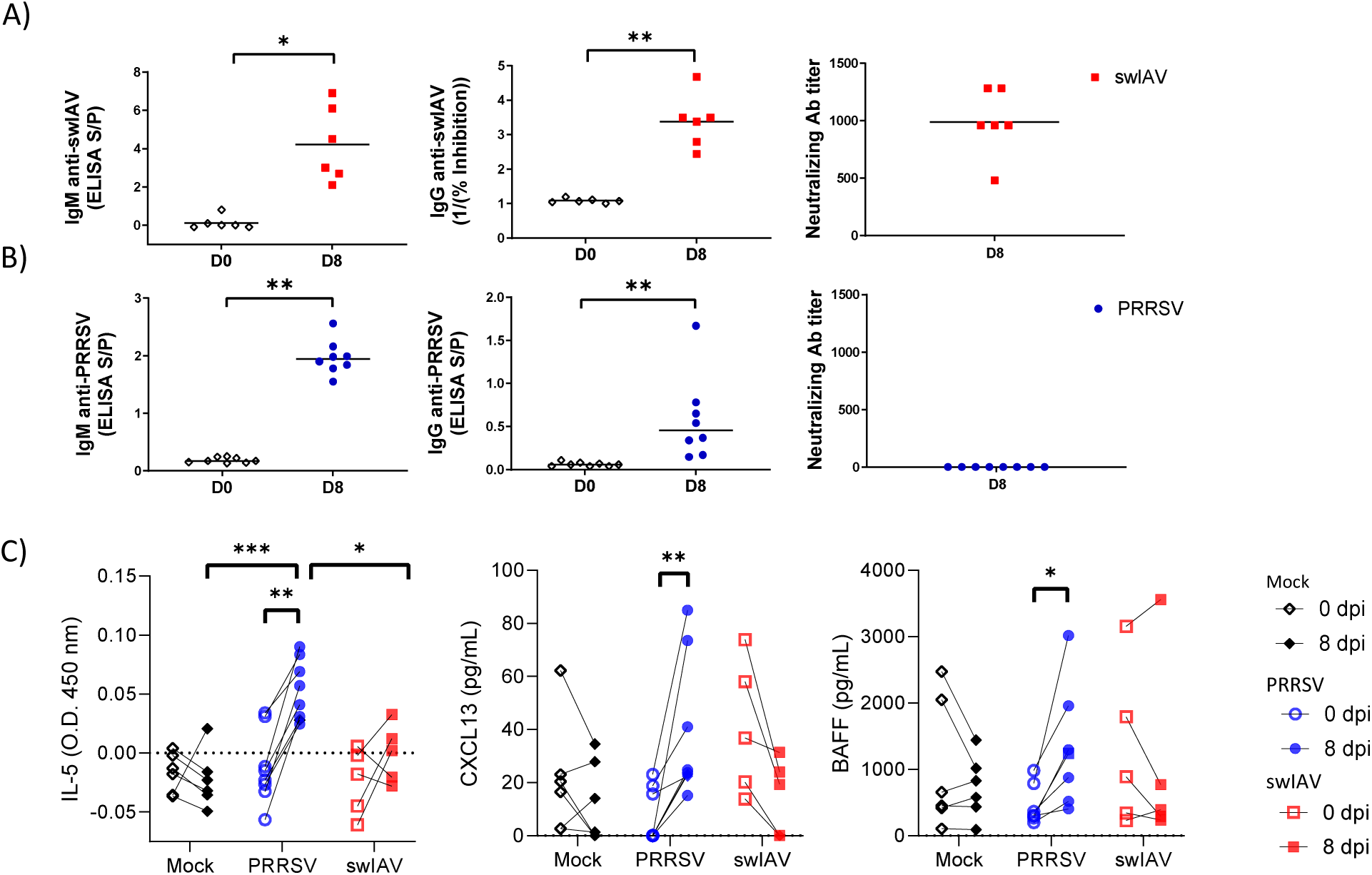
Anti-PRRSV and swIAV serum antibodies, neutralization and B cell maturation related cytokines. Pigs were exposed to either PBS (Mock - black - n=6), PRRSV Lena strain (PRRSV - blue - n=8) or swIAV H1N2 strain (swIAV - red - n=6). Serum was sampled at 0 (empty) and 8 (full) days post-infection. Individual IgM, IgG and neutralizing antibodies titers against swIAV (A) or PRRSV (B). IgM levels are represented as S/P ratio. IgG levels are represented as 1/(inhibition percentage). Neutralization titers corresponded to the reciprocal of the highest dilution of serum able to neutralize the virus. Bars represent means. C) IL-5, CXCL13 and BAFF concentrations in serum. Wilcoxon matched-pairs signed rank test, *p* values: *<0.05, **<0.01, ***0.001.

At 0 and 8 dpi, different cytokines and chemokines known to have roles in the B cell differentiation and maturation processes were titrated. No difference between infectious conditions and time points were observed for IL-2, IL-4, IL-6, IL-10, IL-13, IL-17 nor IFN-γ (data not shown). IL-5 concentrations at 8 dpi were significantly higher in PRRSV-1 compared to Mock- and swIAV-infected animals and presented a significant increase between 0 and 8 dpi (**Fig. 2C**). Two other cytokines, CXCL13 and BAFF, presented a similar significant increase between 0 and 8 dpi in the PRRSV-infected group but not in the Mock and swIAV groups (**Fig. 2C**).

### Upon PRRSV Lena infection, centrocytes were reduced in number and presented an perifollicular location

Interestingly, and despite higher viral loads and clinical scores, LN cellularity at culling (8 dpi) indicated a lower intranodal reaction upon PRRSV compared with swIAV infection (**Supp. Fig. 1A**). To note, one swIAV-infected animal has been excluded from the analysis because of its abnormally fast settlement of the influenza infection, with no more virus shedding as soon as 6 dpi (**Supp. Fig. 1B**), associated with a LN cellular count similar to that of the Mock-infected animals at 8 dpi (**Supp. Fig. 1C**).

After LN cell counting, we stained, analyzed and sorted LN MΦ and B cell at different maturation stages by flow cytometry as previously described (19). Namely, CD169^pos^/CD21^pos^/IgM^neg^ **CD169^pos^ CB**, CD169^neg^/CD21^pos^/IgM^neg^ **CB**, CD21^low^/IgM^pos^/FSC^low^ **CC**, CD21^high^/IgM^pos^ FSC^high^ plasmablasts (**PB**) and CD21^neg^/IgM^pos^/CD172a^neg^/FSC^high^ plasmacells (**PC**) were sorted (**Supp. Fig. 2**). CB sorting was validated on Mock-infected animals using RT-qPCR by the significant over-expression of topoisomerase IIa (Topo2a), a marker of proliferating cells, and of activation-induced cytidine deaminase (AICDA), an enzyme involved in somatic hypermutations (SHM) (**Fig. 3A**). Although non-significant, the expression of Bcl6, a master transcription factor of CB, was higher in CB compared with PB and PC, as expected. Similarly, Blimp1, a marker of terminally differentiated B cells, presented a slightly higher expression in PC (**Fig. 3A**). CXCR5 is the receptor for CXCL13, a chemokine expressed by follicular Dendritic Cells (fDC) and responsible for B cell recruitment in the B cell follicle (28). CXCR5 presented no differential expression among B cell populations of Mock-infected animal LN (**Fig. 3A**).

**Figure 3:**
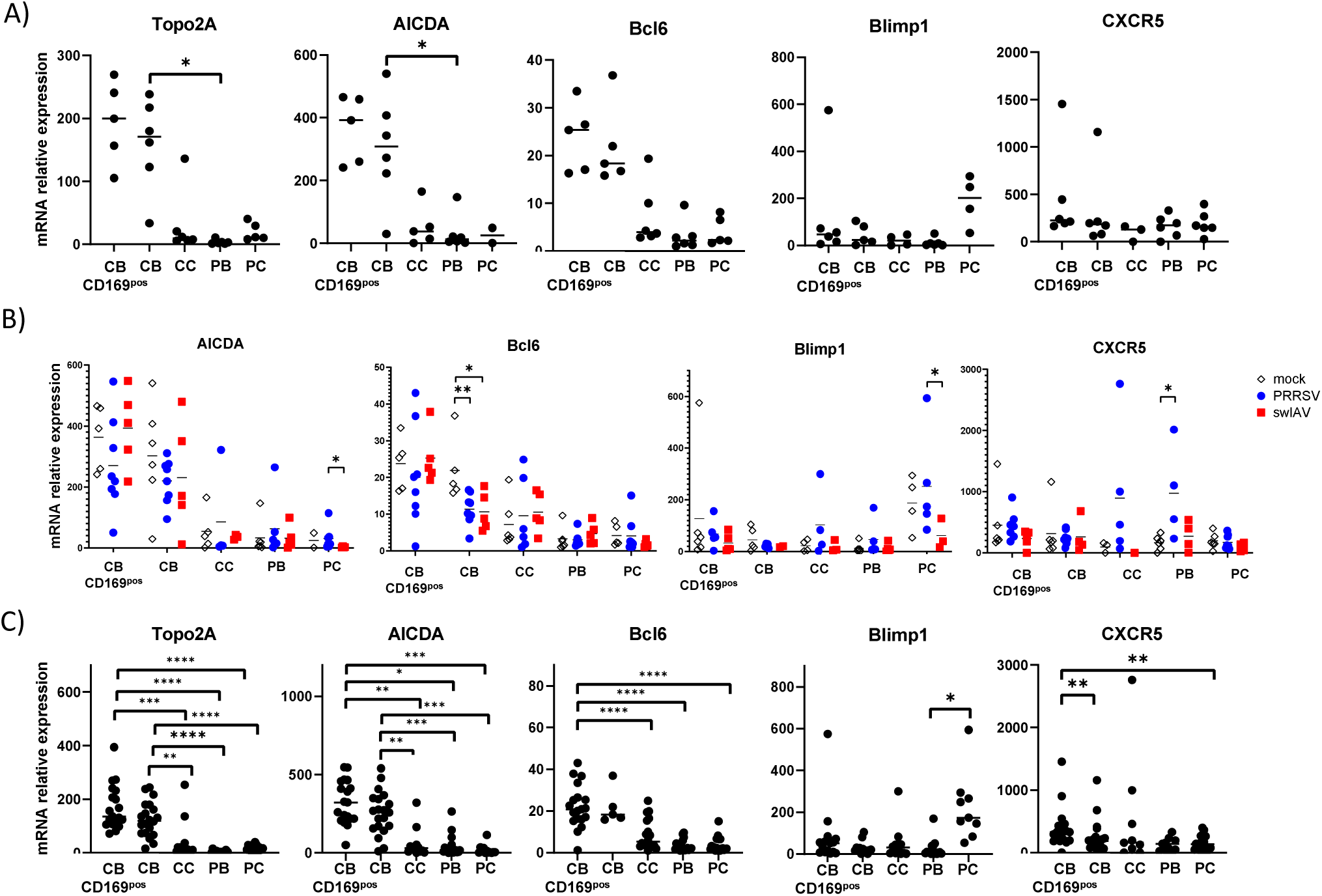
B cells maturation stages specific gene expressions. Lymph node cells were stained for macrophage subpopulations phenotyping, analyzed (see **Sup. Fig. 2**) and sorted using a flow cytometer cell sorter. The relative expression of genes specific of different stages of B cell maturation were measured by RT-qPCR. **A**) B cell maturation stages sorted from Mock-infected animals LN. B) Comparative transcriptomic expression of B cell maturation steps according to infectious status (Mock - black, PRRSV - blue, swIAV - red). C) Comparative transcriptomic expression of B cell maturation steps with pooled infectious conditions whenever no gene expressions differences were observed in **B**). Bars represent means. A) and C) Wilcoxon matched-pairs signed rank test, B) Mann-Whitney unpaired two-tailed t-tests, *p*- value: *<0.05, **<0.01, ***<0.001. CB CD169^pos^: CD169-positive centroblasts, CB: CD169-negative centroblasts, CC: centrocytes, PB: plasmablasts, PC: plasmacells, AICDA: activation-induced cytidine deaminase, Topo2a: topoisomerase IIa.

Upon infection, we observed a higher expression of AICDA and Blimp1 in PC from PRRSV- compared with swIAV-infected animals, a lower expression of Bcl6 in CB of PRRSV- and swIAV- compared with Mock-infected animals, as well as a higher CXCR5 expression in PB from PRRSV- compared with Mock-infected animals (**Fig. 3B**). Topo2a presented no significant difference among the infectious conditions (data not shown).

To increase the statistical power for comparing gene expression in the different B cell populations, we pooled the data from each population (CD169^pos^ CB, CB, CC, PB, PC) from the different infectious statuses after having excluded the populations in which markers presented significant differences compared to Mock: Bcl6 expressions in swIAV and PRRSV CB, Blimp1 in swIAV PC and CXCR5 in PRRSV PB (**Fig. 3B**). We then observed a significant over-expression of Topo2A, AICDA and Bcl6 in CB (both CD169^pos^ and CD169^neg^) as well as an over-expression of Blimp1 in PC compared with all other B cell populations, validating further our cell sorting strategy. Interestingly, we also found an over- expression of CXCR5 in CD169^pos^CB compared with regular CB (**Fig. 3C**).

Having validated our sorting strategy, we then enumerated the different B cell populations. As expected by the higher cellularity in LN from swIAV-infected animals, all the B cell populations (CD169^pos^ CB, CB, CC and PB) were over-represented in swIAV-infected compared with Mock- and/or PRRSV-infected animals, with the striking exception of PC, which appeared as numerous in PRRSV-infected as in swIAV-infected animals LN (**Fig. 4A**). We then performed LN tissue immunofluorescent staining (**Supp. Fig. 3**) in order to analyze the intra- or extra-follicular positioning of the different B cell maturation steps. LN from Mock- and swIAV-infected animals offered a classical morphology. However, PRRSV-infected animals LN systematically displayed a disturbed effMΦ area, associated with a lower conservation of the whole LN parenchymal structure (**Supp. Fig. 3**). Despite this difficulty, we were able to localize CD21^pos^ follicles and to count Bcl6^pos^ CB, and Pax5^pos^ CB and CC inside and outside the follicles. Bcl6^pos^ cells were almost exclusively intrafollicular in all conditions (perifollicular Bcl6^pos^ cells: Mock 1%+/- 2%, PRRSV 9%+/-9%, swIAV 2%+/-3%). Pax5^pos^ cells were also mostly perifollicular. However, PRRSV-infected animals presented significantly more perifollicular Pax5^pos^ B cells than swIAV-infected animals (perifollicular Pax5^pos^ cells: Mock 13%+/-11%, PRRSV 23%+/-7%, swIAV 16%+/-7%) (**Fig. 4B**).

**Figure 4:**
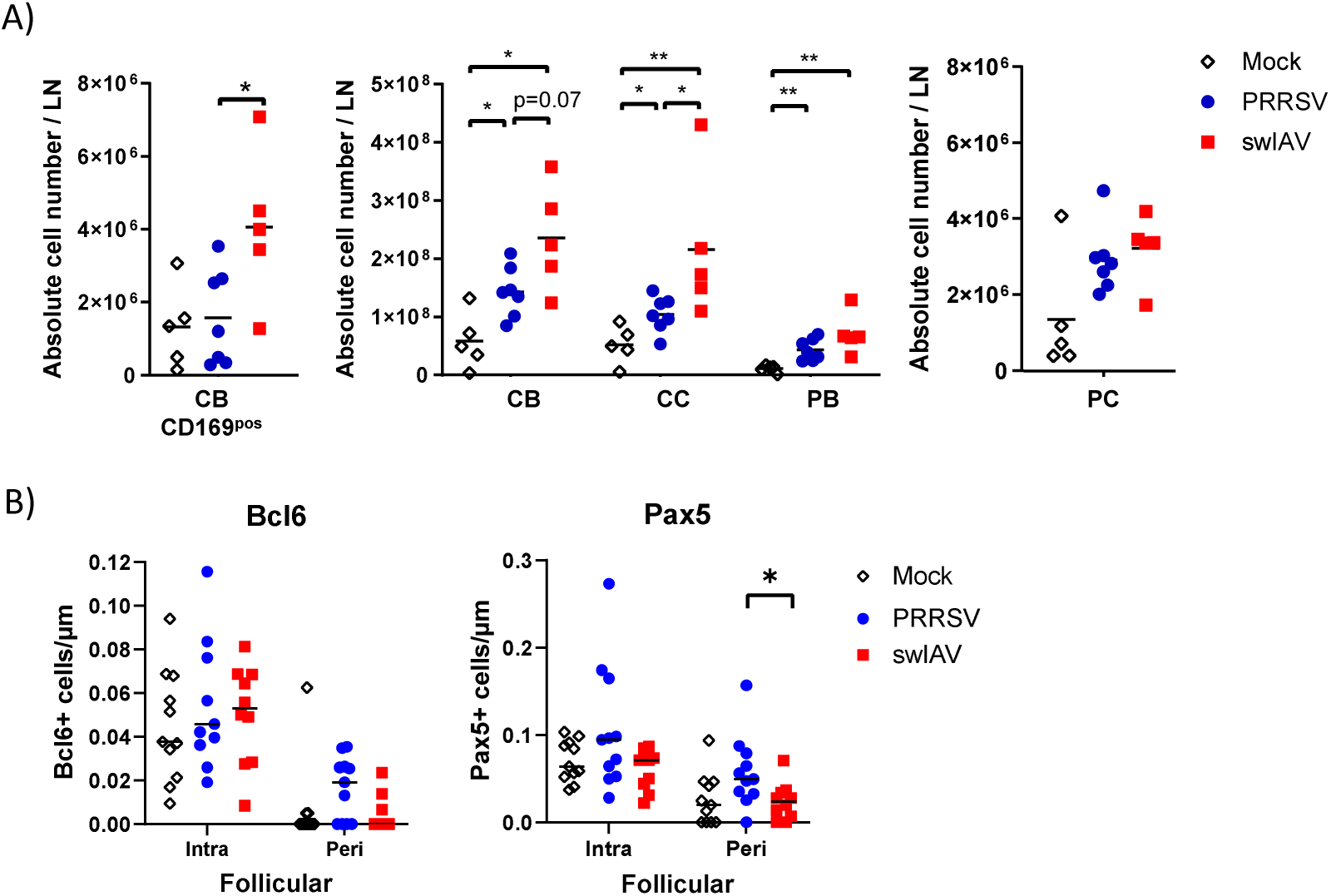
LN B cells subpopulations count in the complete LN or at specific locations at day 8 post PRRSV-Lena or swIAV-H1N2 infections. A) Absolute B lymphocyte LN numbers at different maturation steps, as calculated from absolute total cell number count (**Supp. Fig.1A**) and B lymphocyte percentages determined by flow cytometry (**Supp. Fig. 2**). B) LN B cells count inthe intra-follicular and perifollicular area obtained by microscopic analysis of whole LN (see **Sup. Fig. 3**) stained for CD21, Bcl6 and Pax5. Bars represent means. A) Wilcoxon matched-pairs signed rank test B) Mann Whitney unpaired two-tailed t-tests, *p*-value: *<0.05, **<0.01.

### PRRSV Lena infection triggered the incoming of inflammatory monocytes in T cell area

LN MΦ were defined as CD163^pos^/CD169^neg^ MΦ, CD163^pos^/CD169^pos^ effMΦ and CD163^neg^/CD169^pos^ pfMΦ. We validated the LN MΦ sorting by measuring the expression of previously tested (19) MΦ differentiation genes : CSF1R and MAFB (**Supp. Fig. 4A**). Six sorted populations did not transcriptionally express MΦ markers’ positive controls and were excluded from further analysis: Mock-infected animal 12 CD163^pos^/CD169^neg^ MΦ and effMΦ, Mock-infected animal 13 pfMΦ; PRRSV-infected animal 11 CD163^pos^/CD169^neg^ MΦ and swIAV-infected animal 7 CD163^pos^/CD169^neg^ MΦ and eff MΦ. We then tested genes potentially involved in MΦ functions, CCR2, IL-6, iNOS, PD-L1, CCL2 and CXCL13, and observed no significant differences between MΦ populations from mock-infected animals (**Supp. Fig. 4B and 4C**). CCL2, a chemokine involved in the attraction of inflammatory cells including blood monocytes (29), presented a tendency to be more expressed in effMΦ compared with CD163pos/CD169neg MΦ and pfMΦ (p=0.06 and p=0.1 respectively) (**Supp. Fig. 4C**).

By combining whole LN cell count with the percentage of each population, as measured by flow cytometry, we were able to calculate an absolute number of cells in each MΦ populations. Contrarily to B cells, the higher cellularity in LN from swIAV-infected animals did not significantly impact the MΦ populations in swIAV- compared with Mock- or PRRSV-infected animals LN (**Fig. 5A**). However, CD163^pos^/CD169^neg^ MΦ were significantly over-represented in LN form PRRSV-infected animals as compared with Mock- (50 times increase) and swIAV- (10 times increase) infected animals (**Fig. 5A**).

**Figure 5:**
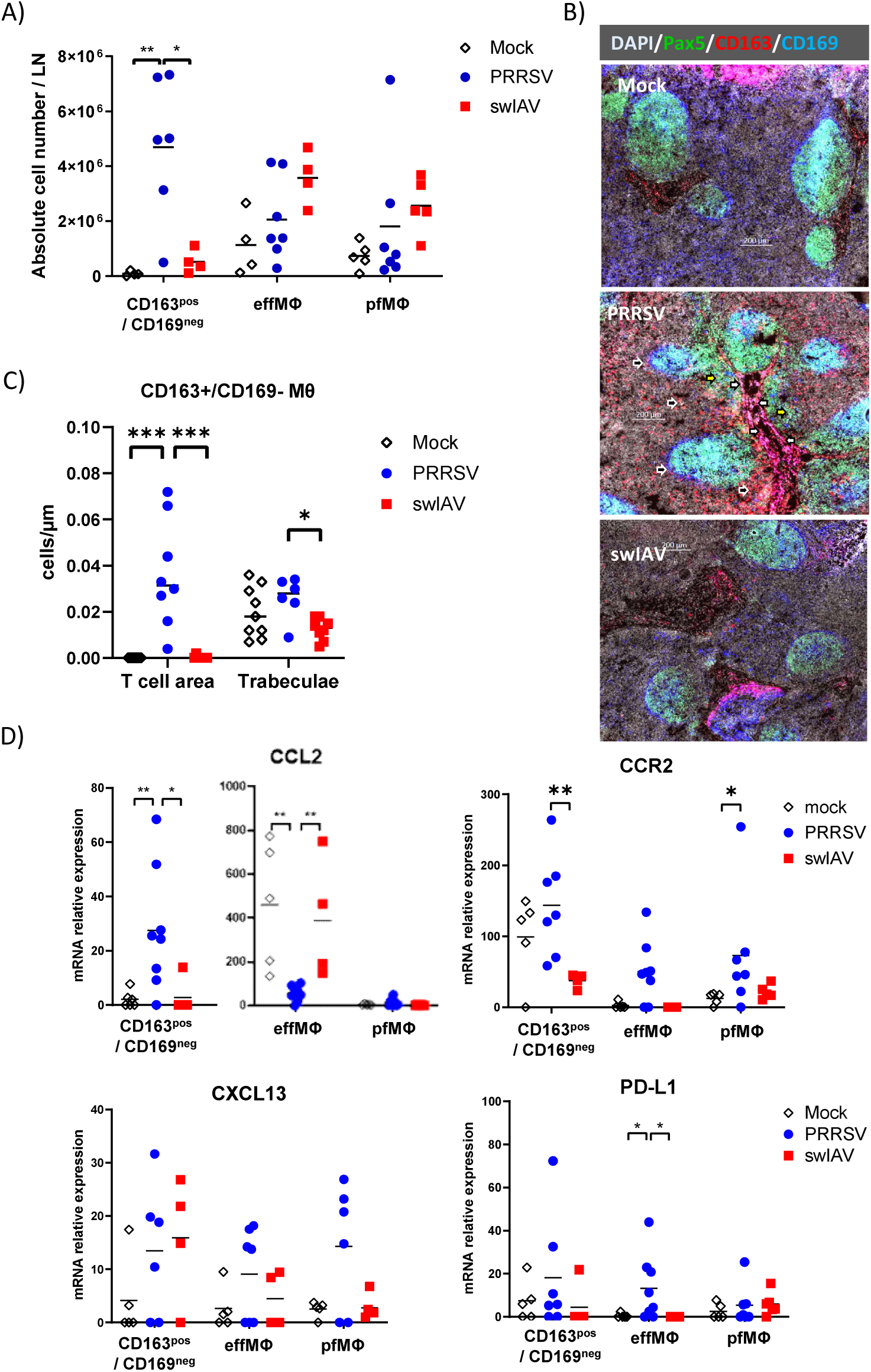
Lymph node macrophages subpopulations count, location and gene expressions at day 8 post PRRSV-Lena or swIAV-H1N2 infections. A) Lymph node cells were stained for macrophage subpopulations phenotyping, analyzed (see **Sup. Fig. 2**) and sorted using a flow cytometer cell sorter. Absolute macrophage subpopulations numbers, as calculated from absolute total cell number count (**Supp. Fig.1A**) and flow cytometry macrophage subpopulations percentages. B) & C) Microscopic analysis of whole LN (see **Sup. Fig. 5**) stained for Pax5 (green), CD163 (red) and CD169 (blue). B) representative images of trachea bronchial LN from each infectious condition. Black arrows with white outlines and white arrows with black outlines indicate CD163^pos^/CD169^neg^ MΦ in the T cell areas and trabeculae, respectively. C) LN macrophage subpopulations count in T cell areas and trabeculae as defined in Material and Methods. D) CCL2 and PD-L1 relative transcriptomic expressions of sorted macrophage subpopulations. CD163^pos^/CD169^neg^: CD163^pos^/CD169^neg^ macrophages, LN: lymph node, effMΦ: efferent macrophages, pfMΦ: perifollicular macrophages. Wilcoxon matched-pairs signed rank test, *p*-value: *<0.05, **<0.01, ***<0.001.

To localize the LN CD163^pos^/CD169^neg^ MΦ appearing upon PRRSV infection, we proceeded to a fluorescent microscopy analysis. B cell follicles were identified through Pax5 staining and the CD163^pos^/CD169^neg^ MΦ population was distinguished through CD163 and CD169 differential expressions (**Supp. Fig. 5**). We localized and counted the CD163^pos^/CD169^neg^ MΦ present in the trabeculae and sinuses (CD163^pos^/CD169^neg^ trabecular MΦ) as well as those located in the T cell area between the B cell follicles (CD163^pos^/CD169^neg^ T cell zone MΦ). Whereas Mock- and swIAV-infected animals presented virtually no CD163^pos^/CD169^neg^ MΦ in T cell areas, PRRSV infected animals presented a strong influx of these cells (**Fig. 5B and 5C**). In addition, swIAV-infected animals presented less CD163^pos^/CD169^neg^ MΦ in the trabeculae and the sinuses than PRRSV-infected animals (**Fig. 5C**). Thus, the CD163^pos^/CD169^neg^ MΦ that appear upon PRRSV-infection are associated with the T cell area rather than the trabecula, and have likely different origins and functions compared to steady state CD163^pos^/CD169^neg^ MΦ. Indeed, RT-qPCR on sorted MΦ populations highlighted the over-expression of CCL2 gene in CD163^pos^/CD169^neg^ MΦ from PRRSV-infected animals compared with Mock- and swIAV-infected animals, in agreement with a pro-inflammatory signature (**Fig. 5D**). Conversely, effMΦ from PRRSV-infected animals strongly down-regulated CCL2 compared with Mock- and swIAV-infected animals. When compared between PRRSV and swIAV conditions, CD163^pos^/CD169^neg^ MΦ presented a higher expression of CCR2, the CCL2 receptor, upon PRRSV infection. CXCL13 expression was not significantly different between MΦ populations or infectious status, although pfMΦ from PRRSV-infected animals presented high CXCL13 expression in 4 out of 6 animals (**Fig. 5D**). Finally, effMΦ from PRRSV-infected animals presented an up-regulation of the immune checkpoint-ligand PD-L1 (**Fig. 5D**).

## Discussion

### Porcine CD169^pos^ centroblasts

First of all, this work allowed us to complete and strengthen our previous results on B cell differentiation steps in the swine inverted LN (19, 20). We confirmed our CB identification strategy as both CD169^pos^ and regular CB presented Topo2A and Bcl6 over-expression, as previously validated (19); moreover, they also presented a clear over-expression of AICDA, an enzyme involved in SHM, a strong marker of CB. In addition, we observed for the first time a differential expression between the two CB sorted clusters: CXCR5 presented a higher expression on CD169^pos^ than on regular CB. CXCR5 is the chemokine receptor involved in the migration toward CXCL13, the chemokine which gradient allows the shuttling of B cells transporting antigens, as described in mice (30–32). We previously showed that CD169^pos^ CB expressed membrane CD169 in the absence of detectable CD169 mRNA, hypothesizing that this CD169 display might result from CB/pfMΦ membrane exchanges during antigen transfer from pfMΦ to CB (19). CXCL13 is secreted by follicular stromal cells (33–35), creating a gradient attracting CXCR5-expressing cells towards the follicle center through complex interactions with EIB2/7α,25-dihydroxycholesterol, sphingosine-1-phosphate (S1P)/S1P-receptor and CXCL12/CXCR4 (for review (36, 37). Thus, CXCR5 over-expression on CD169^pos^ CB reinforces our hypothesis of a role of these cells in transporting free antigens from pfMΦ (at the follicle periphery) to fDC (in the follicle light zone) (30–32).

### Serum cytokines and extrafollicular centrocytes

CXCL13 and IL-5 serum levels have been investigated during HIV chronic infections in human. It has been reported by two groups that CXCL13 serum concentration was positively correlated with NAb responses (38, 39). Conversely, in two other studies, CXCL13 appeared overexpressed in patients with progressive disease compare with elite controller (HIV-infected patients who naturally controlled their viral load) and patients on antiretroviral therapy (40, 41). In this last study, contrarily to CXCL13, IL-5 correlated with nAb breadth (41). In a PRRSV context, it seems that IL-5, CXCL13 and BAFF, are the markers of an unresolved immune response trying to unsuccessfully control the viral load rather than markers of an efficient neutralizing immune response. Accordingly, it would be of great interest to compare the serum expression of these cytokines and chemokines on a daily basis during the early steps of swIAV and PRRSV infections in pigs.

Despite ongoing viral shedding and a higher clinical score, we observed lower total LN cellularity in PRRSV- compared to swIAV-infected animals LN. Notably, this deficit was paralleled by CB, CC and PB, but not PC, decreases. One possible explanation for this discrepancy is an extrafollicular differentiation of PC during PRRSV infection. To investigate this possibility, we checked for extrafollicular B cell differentiation and observed an increase in perifollicular Pax5^pos^ B cells upon PRRSV infection. Whether these cells account for extrafollicular B cell differentiation that would skip the intermediate follicular steps and the ensuing Ab affinity maturation (for review (42)), as postulated by others (15, 16), remains to be explored in more detail.

Concerning B lymphocytes maturation steps, the only gene expression differences between PRRSV- and swIAV-infected animals were the over-expression of AICDA and Blimp1 in PC. AICDA expression in PC usually reflects isotype switching, which might again be the consequence of the ongoing PRRSV immune response, as might be the Blimp1 overexpression, compared with the resolving anti-swIAV infection. However, the main role of AICDA is to trigger SHM, and SHM occurring in PC have been associated with extrafollicular B cells maturation (43, 44). Thus, the over-expression of AICDA in PC is in agreement with extrafollicular B cells maturation.

### Inflammatory monocytes/macrophages and PD-L1

Upon PRRSV infection, we observed an influx of CD163^pos^/CD169^neg^ monocytes/MΦ in the LN T cell area. According to murine data, at steady state, this MΦ phenotype might include different LN populations, such as medullary cord MΦ and T cell zone MΦ (for review (45)).

In the murine model of acute and chronic lymphocytic choriomeningitis virus (LCMV), it has been shown that chronic strains were endowed with the unique capacity to attract inflammatory myeloid cells which inhibit both T (46) and B cell (47) mediated immune responses. In agreement with this model, we observed an increase of CCL2 expression in CD163^pos^/CD169^neg^ MΦ in LN of PRRSV-infected pigs, consistent with an inflammatory monocyte signature. Interestingly, an influx of uncharacterized MΦ in the T cell zone of pig LN has already been observed with different PRRSV strains, although the pattern was more pronounced with the highly pathogenic SU1-Bel strain (48). According that the Lena strain used here was also highly pathogenic, a relation between pathogeny and LN MΦ influx in the LN can be proposed. Furthermore, while the CD163^pos^ MΦ population increases and up-regulates CCL2 upon PRRSV infection, effMΦ strongly down-regulate CCL2 expression. This might lead to the weakening of the CCL2 gradient present in Mock- and swIAV- infected animals, from the T cell area toward the efferent area. The role of this LN gradient in normal and altered immune responses remains to be explored.

PD-1, the PD-L1 receptor, is expressed on B cells and acts as an inhibitor of the B cell activation cascade (49). PD-L1 has been shown to be over-expressed in the thymus (27), broncho-alveolar lavage (50), lung tissue, in *in vivo* infected AM (51) as well as in LN (52) of animals infected with different strains of PRRSV-1. We observed a PD-L1 over-expression in effMΦ, the porcine counterpart of medullary MΦ (19), which are thought to aid in the final maturation of PB into PC in the murine model (53). Indeed, the efferent area is the porcine LN region where mature or maturing Blimp-1^pos^ B cells accumulate in close proximity to effMΦ (20). Anti-porcine PD-L1 nAb have been developed (54) and would be worth testing in PRRSV-infectious settings.

### Conclusion

Herein, we were able to investigate in detail the differential impact of swIAV and PRRSV infections on B cell maturation. By this PRRSV/swIAV comparison in growing, conventional animals, we comforted two of the main hypotheses previously proposed for both PRRSV-1 and PRRSV-2 independently of pathogenicity level: the likely induction of extrafollicular B cell activation (16, 17) and the over- expression of PD-L1 in different tissue MΦ (27, 50, 52). We also evidenced the invasion of LN T cell areas of PRRSV-infected animals by inflammatory monocytes/MΦ that may have a strong impact on the B cell maturation process, as described for LCMV in mice (47). Although this last feature might be related to the level of PRRSV pathogenicity. Attenuated vaccines are one of the main resources to control PRRSV in farms. However, they still have similar abilities to inhibit the immune system as wild-type PRRSV, which contributes to their low efficacy and raise concerns about their use (55). PRRSV inhibition of the B cell response induction seems multifactorial and it remains to be explored which PRRSV-genes/factors are involved in these different aspects of the immune blocking, in order to design efficient attenuated vaccines depleted of these properties.

## Authors contributions

CH prepared the viruses for inoculation. CH, MM and NB proceeded the serum, lung and lymph node tissues. JA and NB performed cell sorting. AP and NB carried out immunostaining of LN slides. CH and AP conducted RT-qPCR. PR, CD, GS and OB handled immunoglobulin titrations and neutralizing antibodies determinations. CH, PR, CD, FM, JA, GS, OB PM and NB corrected and edited the manuscript, providing thorough discussions and critical revisions. NB supervised the work, designed the experiments, analyzed the images, prepared the figures and wrote the manuscript. All authors contributed to the article and approved the submitted version.

## Acknowledgments

The authors would like to thank Mireille Ledevin and Laurence Dubreil from the Apex facility (Nantes, France); Alfred Bensaid, JI Nunez, Joachim Segales from CRESA (Barcelona, Spain) ; Jorge Díaz Pedroza from CMCIB/IGTP (Barcelona, Spain) and Roselyne Fonseca Sanchez-Gesny from ANSES (Ploufragan, France).

