## Supplementary material for "Comparative impact of porcine reproductive and respiratory virus and swine influenza A virus infections on respiratory lymph nodes B cells and macrophages": Supp Data & Table

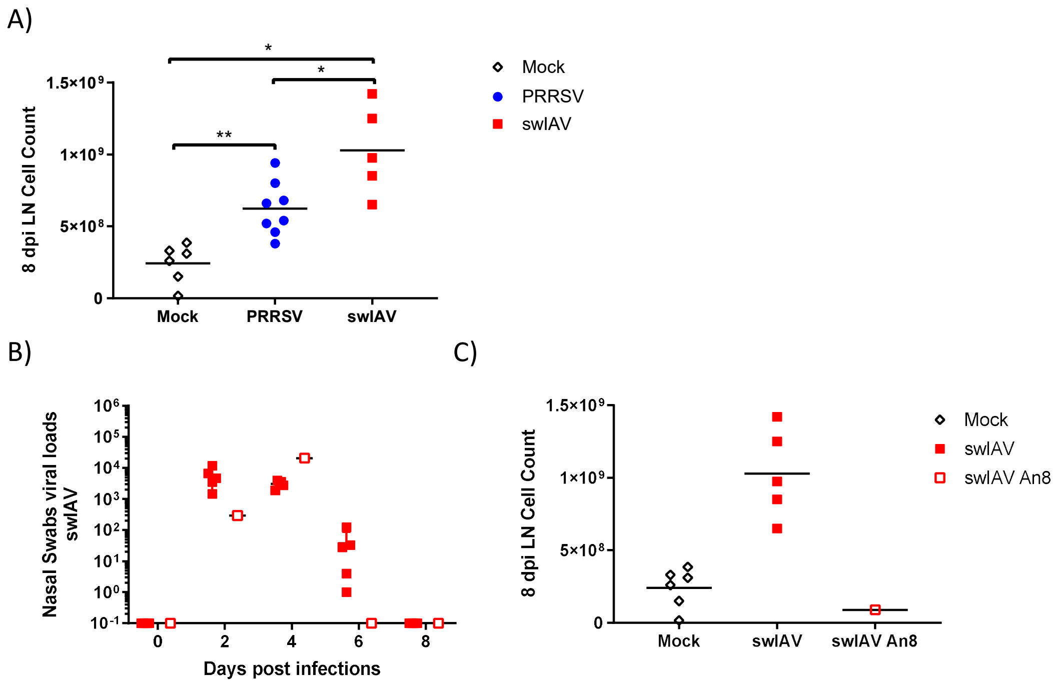


**Supplementary Figure 1**: Total LN cellularity and outlier swIAV-H1N2-infected animal that cleared the virus at 6 dpi and restored normal LN cellularity at 8 dpi.
Total tracheobronchial LN cellularity at 8 dpi A) according to infectious status, B) Nasal swabs were processed every other day and viral load were measured by RT-qPCR. Mock: 6, controlled non-infected animals. swIAV: 5 out of 6 swIAV-H1N2 infected animals. C) Total tracheobronchial LN cellularity at 8 dpi according to animal number 8 swIAV An8: outlier animal 8 swIAV infected animal.

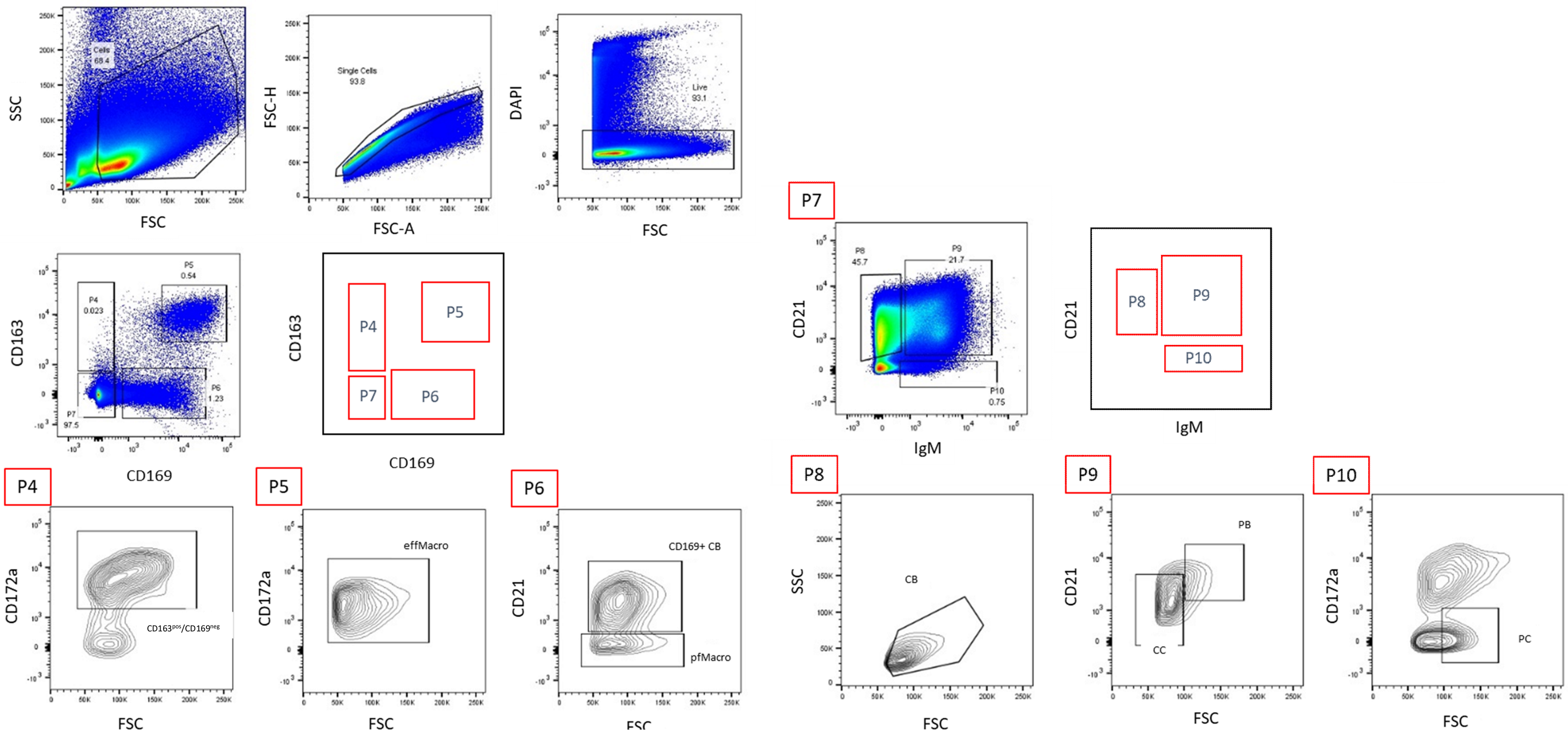


**Supplementary Figure 2**: Flow cytometry gating strategy for LN Macrophages and B cells analysis and sorting.
DAPI, CD163-PE, CD169-PE-Cy7, CD21-FITC, IgM-Alexa-647 and CD172a-PE-Cy7 staining allow FSC/SCSC followed by singlet live gating.
Then CD163/CD169 depicting allowed the gating of the populations P4, P5, P6 and P7. P4 and P5 gated-populations were re-gated on CD172a/FSC dotplot order to sort respectively CD163$^{pos}$/CD169$^{neg}$ macrophages and efferent macrophages (effMΦ). P6 gated-population was re-gated on CD21 and FSC dotplot in order to sort CD21-negative perifollicular macrophages (pfMΦ), and CD21-positive centroblasts (CD169$^{pos}$ CB). P7 population was regated on CD21/IgM dotplot to gate CD21-positive/IgM-negative P8, CD21-posititive/IgM-positive P9 and CD21-negative/IgM-positive P10 populations. P8 population, regated on FSC/SSC large cells were sorted as centroblasts (CB). P9 populations regated on CD21/FSC dotplot were sorted as CD21-low/FSC-low centrocytes (CC) and CD21-high/FSC-high palsmablasts (PB). Finally P10 population was sorted as CD172a-negative/FSC-high plasmacells (PC).

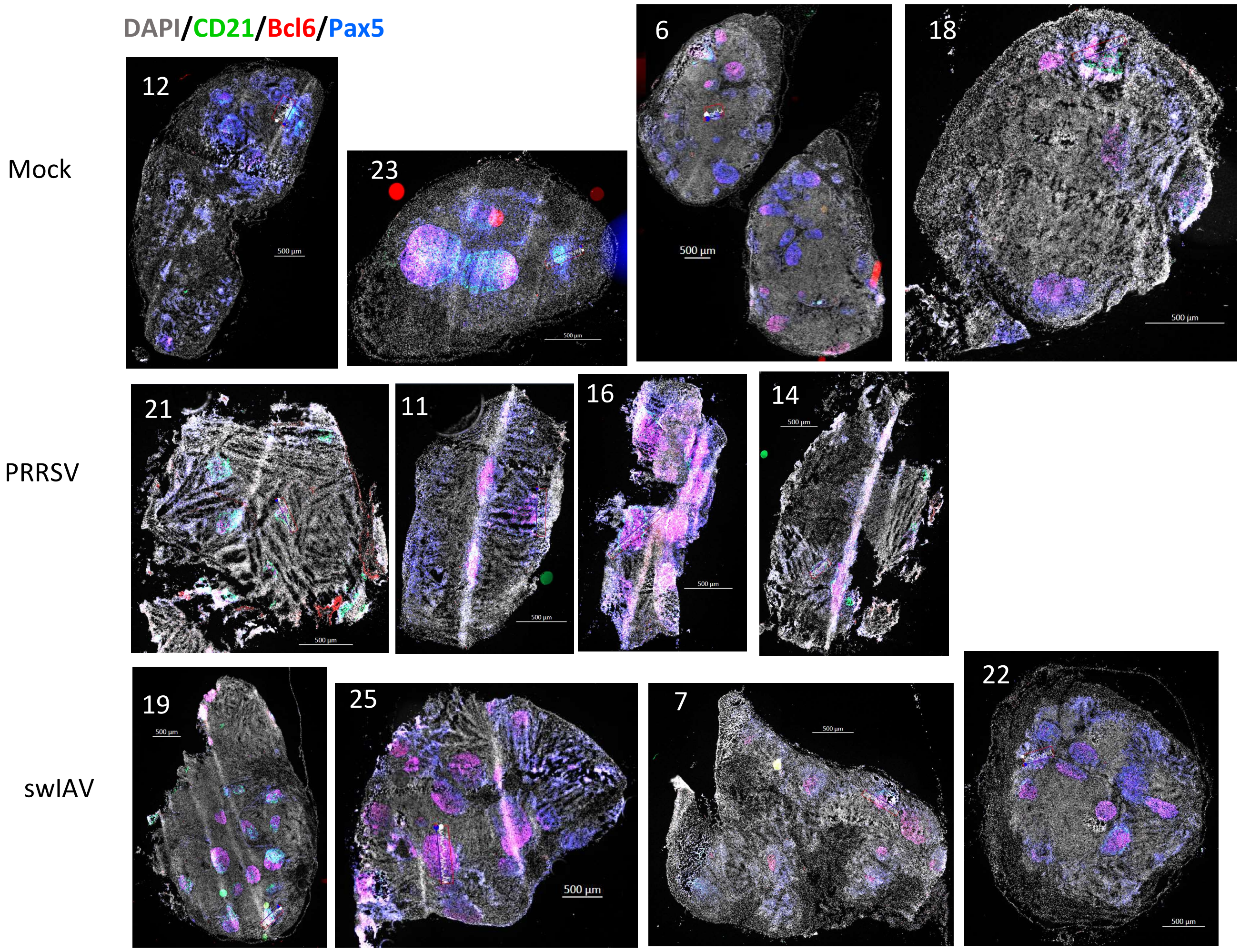


**Supplementary Figure 3**: DAPI/CD21-A488, Bcl6-A555 and Pax5-A647 staining for CB and CC counting using whole lymph nodes microscopic analysis according to their follicular or extrafollicular location. Numbers on each image is the animal identity number. In each picture, one example of the selected line (between 2 open squares) encompassing a follicle and its perifollicular area (as defined in Mat & Med) is depicted

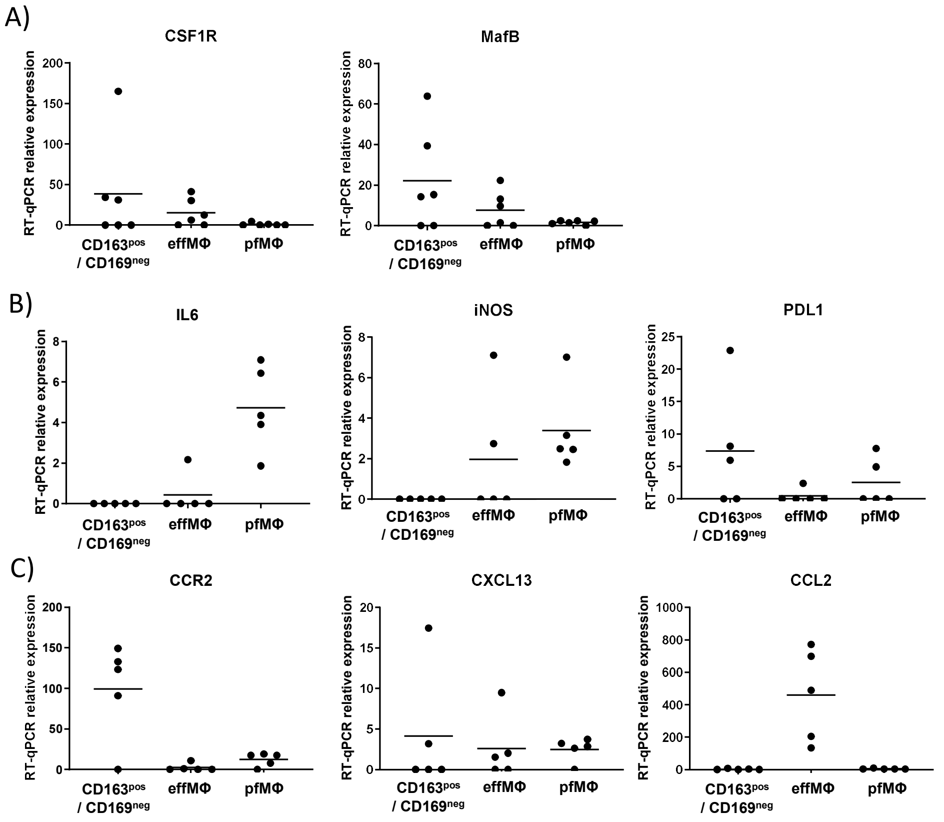


**Supplementary Figure 4**: Sorted LN Macrophage populations gene expression signatures from Mock-infected animals.

RT-qPCR-relative transcriptomic expressions.

CD163$^{pos}$/CD169$^{neg}$: CD163$^{pos}$/CD169$^{neg}$ macrophages. effMΦ: efferent macrophages, pfMΦ: perifollicular macrophages.

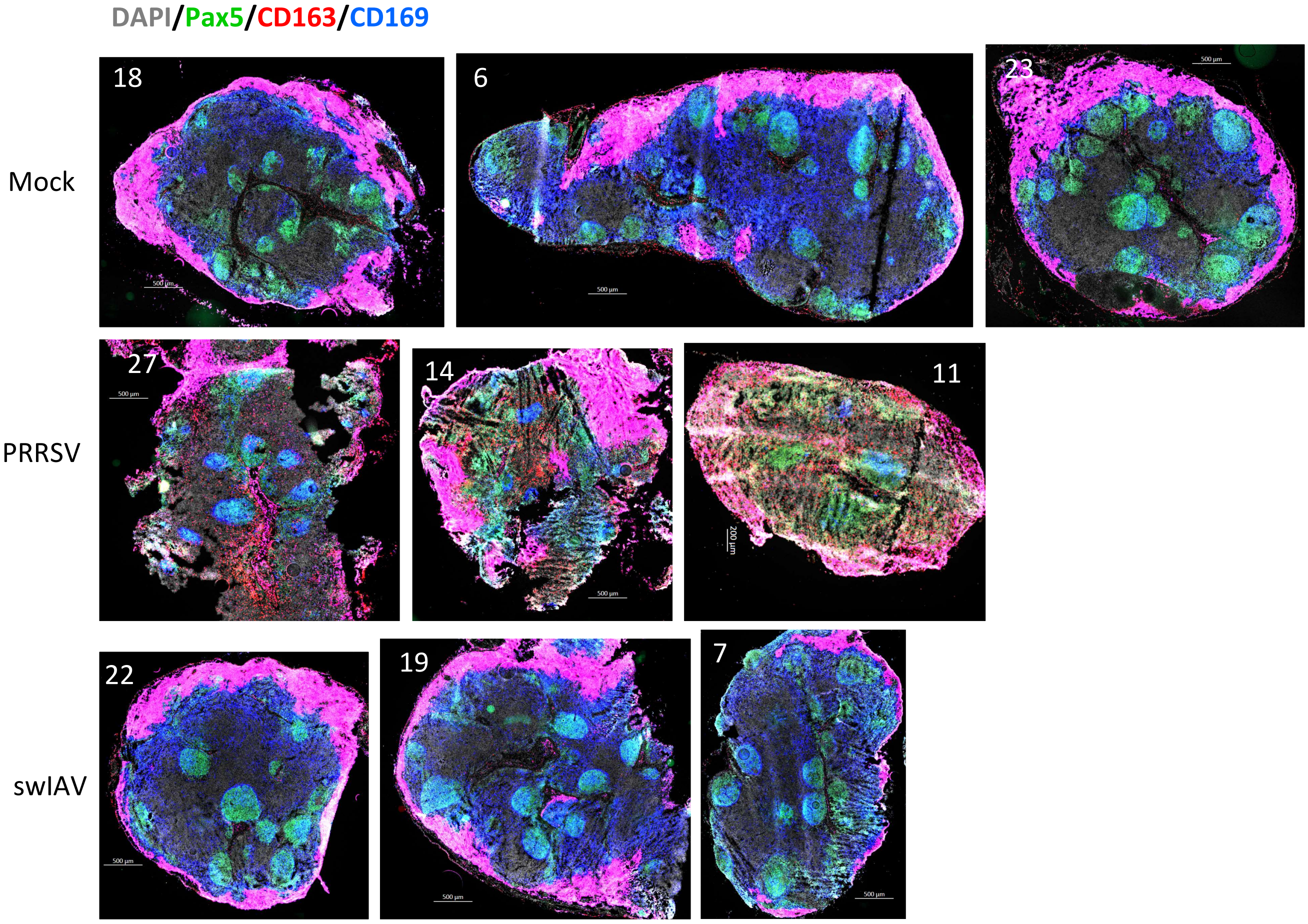


**Supplementary Figure 5**: DAPI/Pax5-A488/CD163-A555/CD169–A647 staining for the positive and negative macrophages counting using whole lymph nodes microscopic analysis. Each picture is identified by the animal identity number.

Supplementary Table 1: Clinical score criterions

| **Variables to consider or score range** | | | **Monitoring frequency** |
|---|---|---|---|
| Rectal Temperature (from 0 to 3) | Normal (38.5 - 40°C) | 0 | Daily |
| | Slightly febrile process (40 – 40.5) | 1 | |
| | Moderate fever (40.5 – 41.5°C) | 2 | |
| | High Fever (> 41.5°C) | 3 | |
| Breathing (from 0 to 3) | Normal | 0 | Daily |
| | Slightly dyspnea | 1 | |
| | Moderate dyspnea | 2 | |
| | Moderate dyspnea with tachypnea | 3 | |
| Behaviour (from 0 to 3) | Active | 0 | Daily |
| | Not too active but responds to the external stimuli | 1 | |
| | Inactive even when stimulated | 2 | |
| | Clearly prostrated | 3 | |
| Bodyweight (from 0 to 3) | Gain weight in a week | 0 | Daily |
| | The animal do not gain weight in the week but there is no significant loss | 1 | |
| | The animal has lost between 2% and 10% of the live weight corresponding to the last weighing | 2 | |
| | Weight loss greater than 10% of the last recorded weight | 3 | |
| **Total score** | | **From 0 to 16** | |
